# Capsicum cultivated under adverse conditions in Southern Japan produces higher concentrations of antioxidants and pungent components

**DOI:** 10.1101/719740

**Authors:** Katsuko Kajiya, Hiroki Yamanouchi, Yurika Tanaka, Hiroka Hayashi, Yuji Minami

## Abstract

Growing crops in sabulous soils is often challenging due to their limited oligotrophy and weak water retention. However, some plants adapt to these adverse growth conditions, and in some cases, favorable properties are imparted to the fruit. This study investigated the influence of the cultivation environment on Capsicum by assessing the levels and functions of both pungent components and antioxidants when cultivated in sandy soils in Southern Japan; these parameters were then compared to those in traditional tropical-origin Capsicum. In seven varieties of Capsicum, the distribution of capsaicin and dihydrocapsaicin differed between the pericarp and seeds within the placenta. The leaves and fruits of Habanero orange and Tabasco, which are the most suitable for cultivation in sandy soil, were collected during the cultivation period and analyzed in terms of their size, color, and pungent component composition. Pungent components were detected in fruits only, and not in leaf or flower samples. In particular, we found that pungent components were generally present within the seeds and placenta. Antioxidant activity and nitric oxide production within human vascular endothelial cells were also evaluated to compare the differences in their functionality. *Satsuma*-Habanero orange cultivated under adverse conditions possessed the highest antioxidant activity. Furthermore, *Satsuma*-Capsicum cultivated under adverse conditions exhibited higher levels of antioxidants than traditional tropical-origin peppers, and induced similar levels of nitric oxide production in the vascular endothelial cells. We concluded that Capsicum cultivated in harsh environments produced beneficial effects such as higher levels of antioxidants and capsaicinoids in seeds and placenta. Moreover, the fruits from these plants could be harvested for a significantly longer period and took longer to spoil than traditional Capsicum; thus, they show merit as a viable commercial crop in Japan.

## Introduction

Sandy soil in coastal areas lacks nutrients and has low moisture retention, severely limiting the number of adaptable crop species that can be grown in this soil. The components and nutritional value of plants vary depending on the cultivated environment. Sandy soil has extremely low clay content and little accumulation of organic matter. Further, this soil possesses good breathability and drainage, but low water retention and natural fertility. Moreover, the soil temperature tends to rise rapidly. Therefore, it is susceptible to drought, and nutrients from added fertilizer are often removed due to leaching. The typical properties of sandy soils are strongly influenced by external environmental factors due to their low physical buffer capacity, such as earthiness, air permeability, water permeability, water retention, soil temperature, sand scattering, chemicals, nutrient sources, nutrient transfer, nutrient absorption, and soil pH, as well as biological factors such as pests and the accumulation of organic matter in the soil. The properties of sandy soil are thus variable and unstable.

Capsicum is a plant belonging to the family *Solanaceae*. The plant height of *Capsicum spp*. ranges from 40–60 cm, and the stem splits into many branches. The leaves are mutually alternated with a long petiole and have an oval shape. After the flowers have bloomed, the plant starts bearing green fruits; inside the fruit is a cavity containing the seeds and placenta (s/p). Depending on the variety, the fruit shape may be rounded or short, and the color may differ. Many *Capsicum spp*. are spicy, and are therefore primarily used as spices, except for some sweet peppers, which are generally mild in flavor [1,2]. The main pungent components of Capsicum are capsaicinoids, including homologs such as capsaicin, dihydrocapsaicin, nordihydrocapsaicin, homocapsaicin, and homodihydrocapsaicin [3–5]. In the current study, we focused on capsaicin and dihydrocapsaicin because these compounds possess strong pungency. Moreover, Capsicum generally contains 60–70% capsaicin and 30–40% dihydrocapsaicin, with other capsaicinoids being present in trace amounts [3–5]. The concentration of pungent components in Capsicum can vary depending on the light intensity and temperature at which the plant is grown, the age of the fruit, and the position of the fruit on the plant [4,6]. The nondestructive identification of pungent component contents is useful for determining the optimum harvest time. However, the fruits are currently empirically harvested because the pungent component contents and color of the fruits are irrelevant. Thus, it is important to understand the changes in capsaicin and dihydrocapsaicin contents over time within the Capsicum fruit. Measuring the levels of pungent components in various parts of the Capsicum plant can reveal whether they can be used as a substitute for Capsicum fruits in food.

Oxidative stress, which is caused by excessive production of active oxygen within the body, leads to obesity and lifestyle-related diseases. Therefore, the antioxidant properties of foods are important [7], and peppers are thought to possess considerable antioxidant activity. Further, nitric oxide (NO) is released from the vascular endothelial cells (VECs) to protect blood vessels by regulating their contraction and relaxation and by preventing thrombus formation due to the attachment of white blood cells and other blood components to the vascular endothelium. Traditional tropical-origin peppers influence the release of NO by endothelial cells [8,9]. However, if VECs are damaged by oxidative stress caused by reactive oxygen species or oxidized low-density lipoproteins, the production of NO is suppressed, increasing the risk of cardiovascular diseases. Thus, improving NO production by VECs is critical for protecting blood vessels.

The distribution of pungent components in pepper varieties cultivated in harsh environments has not yet been investigated. This study measured the concentrations of pungent components in the pericarp and placental seeds of *Satsuma*-Capsicum plants cultivated in harsh environments compared to those in traditional topical-origin peppers. We further investigated their antioxidant activity and effect on vascular endothelial function by analyzing the changes in nitric oxide (NO) production in VECs [10].

## Material and Methods

### Chemicals

Acetonitrile and capsaicinoids were purchased from Sigma Aldrich (St. Louis, MO, USA). Ethanol, hexane, dichloromethane, and 2,2-diphenyl-1-picrylhydrazyl (DPPH) were obtained from Fujifilm Wako Chemical Corporation (Osaka, Japan). 2,3-diaminonaphthalene (DAN) was purchased from Dojindo Laboratories (Kumamoto, Japan). Normal human VECs were obtained from Cosmo Bio Co., Ltd. (Tokyo, Japan).

### Sample preparation

A total of seven *Capsicum* species and strains (Figure 1A) were included in this study, including *C. chinense* (Habanero orange and Habanero red), *C. annuum* (Indonesian origin and Laris), and *C. frutescens* (Taruna pepper, Okinawan chili pepper, and Tabasco pepper). The plants were cultivated in sandy soils in southern Japan, and they were named *Satsuma*-Capsicum for convenience. *Satsuma*-Capsicum plants from each of the seven varieties were harvested when the coloration of the mature fruits exceeded 90%; this decision was based on color changes due to differences in maturation time. Two traditional tropical-origin peppers, Habanero orange and Tabasco peppers originating from the USA, were provided by Dr. Jun-Ichi Sakagami and Mr. Kenta Komori at the Laboratory of Tropical Crop Science, Faculty of Agriculture, Kagoshima University (Japan). The peppers were carefully separated into the pericarp and s/p (Figure 1B), then placed into a glass petri dish and dried using a forced convection drying oven (DO-60FA, AS ONE, Osaka, Japan) at 60 °C for 13 h. Thereafter, the thoroughly dried material was crushed using a mortar and pestle.

**Figure 1.**
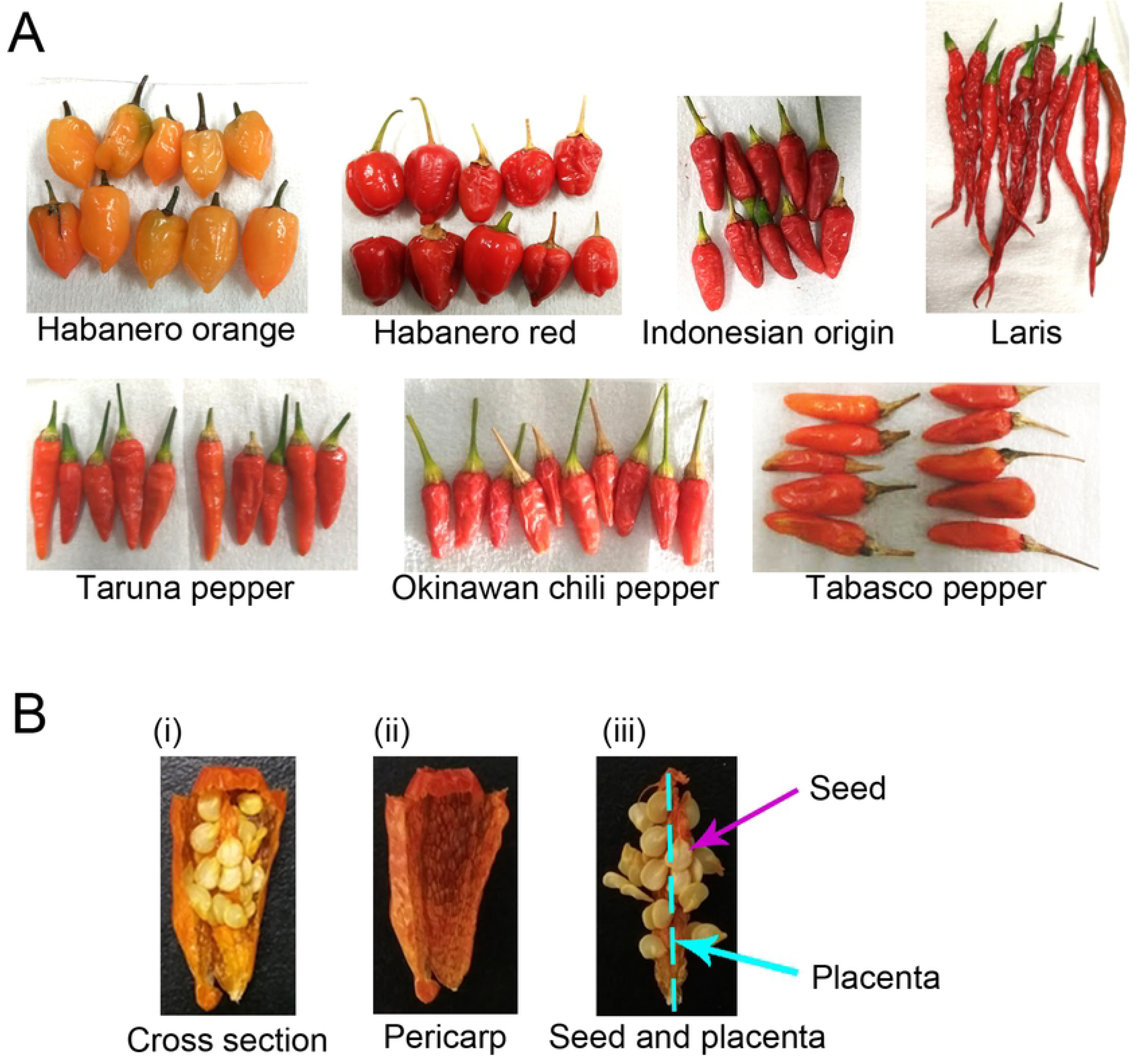
(A) Representative photographs of the seven *Satsuma*-Capsicum strains used in this study. (B) Sample preparation. Peppers were carefully (i) cut vertically, and then separated into (ii) the pericarp and (iii) seeds and placentas.

### Quantitative analysis of pungent components

Each dried sample (10 mg) was added to a microfuge tube containing 1 mL acetonitrile, ultrasonically processed (ASU-6M, AS ONE) for 1 h, and centrifuged at 1,600 × *g* for 10 min at 4 °C; the resulting supernatant was used to measure the levels of pungent compounds. After filtration through a 0.45-μm filter (Toyo Roshi Kaisha, Tokyo Japan), the samples were analyzed using high-pressure liquid chromatography (HPLC; Extrema, Jasco, Tokyo, Japan). The identification of individual compounds was performed using highly selective spectral data according to the retention times of each individual compound on an ultraviolet-photodiode array detector (UV-4075 and MD4010, Jasco). The conditions for HPLC analysis were as follows: the C18 reverse phase column (TSK-gel ODS-100Z, 5 μm, 4.6 mm I.D. × 150 mm, Tosoh, Tokyo, Japan) and guard column (TSK-gel guardgel ODS-100Z, 5 μm, 3.2 mm I.D. × 15 mm, Tosoh) were maintained at 40 °C, and detection was performed at 280 nm. The mobile phase contained 1% acetic acid in water (A) and acetonitrile (B). We used a gradient of 0 min with 50% solution B, and 0–18 min with a direct increase in solution B of up to 75%. The flow rate was 1.0 mL/min, absorption was measured at 280 nm, and the injection volume was 10 μL.

### Measurement of changes in of pungent component contents in leaves and fruits over time

We selected two commonly consumed varieties of *Satsuma*-Capsicum and examined the changes in the levels of pungent components in the leaves and fruits over time. We divided the *Satsuma*-Capsicum tree into the top and bottom sections, and collected three leaves and fruits from each section. Samples were gathered from four *Satsuma*-Capsicum trees. Habanero orange and Tabasco peppers were sowed between April 14th and April 15th, 2017; the peppers were then planted in a sandy soil field on June 16th, 2017. Following planting, leaves were collected 1–2 times per week a total of thirty-two times. Collection started on July 15th (30 days after planting), as soon as the plants showed sufficient growth to collect the leaves without preventing further growth. Collection continued until December 23rd, after which withering and leaf dropping rendered collection impossible. Flowers (harvested between August 10th and 26th, 2017) and fruits (harvested between August 12th and December 23rd, 2017) were also sampled and the levels of pungent components were measured. In order to eliminate individual differences arising from harvesting, at least three leaves were collected from the top and bottom of each plant. Flowers and fruits were also collected in the same way. Prior to harvesting, the overall condition of the plants was measured to ascertain the state of growth. After acquiring images using a standard digital camera (SX720 HS, Canon, Tokyo, Japan), the length, width, and weight were recorded, and the pigment colors were measured with a color-difference meter (CR-20, Konica Minolta, Tokyo, Japan) using four measurement points per sample. Leaf pigment color was expressed using a color code conversion tool (freeware Color Converter, W3Schools) to convert from the L*a*b system to the RGB color coordinate system. L* expresses brightness, where values closer to 0 are closer to white, and values closer to 100 are closer to black. For a*, negative values indicate a shift towards green, and positive values indicate a shift towards red. For b*, negative values indicate a shift towards blue, and positive values indicate a shift towards yellow.

All samples were then cut with scissors. Each sample (1 g) was added to a microfuge tube containing 1 mL ethanol, and incubated at 4 °C for 1 week. The extracted solution from each sample was subjected to ultrasonic processing for 10 min and centrifuged at 1,600 × *g* for 10 min at 4 °C. The resulting supernatant was passed through a 0.45-μm filter and used for measurement. The conditions for HPLC analysis were the same as those for the quantification of pungent components.

### Measurement of antioxidant activity

We used the DPPH method to measure antioxidant activity [11–13]. In brief, each dried sample (50 mg) was placed in a microcentrifuge tube, to which 2 mL hexane/dichloromethane (1:1) solution was added. The mixture was vortexed, ultrasonically treated for 10 min, and centrifuged at 1,600 ×*g* at 4 °C for 10 min; thereafter, 1 mL supernatant was collected and prepared by vacuum concentration (VEC-260, Iwaki, Tokyo, Japan). To the concentrated sample, 1 mL 50% ethanol was added, vortexed, and ultrasonically treated until the concentrate was dissolved. This sample solution (50 μL) was added to each well of a 96-well microplate, and then 50% ethanol was also added into each well as necessary. Then, 50 μL 800 μM DPPH solution was added to the sample solution in the dark and incubated at room temperature (25 °C) for 20 min in the dark. Absorbance was measured at 540 nm via a microplate reader (Infinite 200 PRO, Tecan, Männedorf, Switzerland). A calibration curve was produced using the reference standard compound Trolox with a correlation coefficient of R^2^ = 0.9983. Each experiment was performed in quadruplicate. The DPPH scavenging effect was calculated via the following equation: DPPH scavenging effect (%) = (A0 − A1)/A0) × 100, where A0 is the absorbance of the control reaction, and A1 is the absorbance of the sample or standard.

### NO quantification using a modified Griess method

Typically, NO_2_^−^ is measured using the Griess method [14–16]. The reduction of NO_2_^−^ mediated by nitrate reductase is used to ensure that the NO_2_^−^ concentration represents the original NO level of a sample. A fluorescence method [17] using DAN is a more recently developed NO_2_^−^ assay with higher sensitivity than that of the Griess method. Because NO_2_^−^ reacts with DAN under acidic conditions to form a fluorescent adduct (naphthalenetriazole), we quantified the product by measuring its fluorescence intensity using a microplate reader. VECs from a human coronary artery were seeded at 5.0 × 10^4^ cells/mL and grown in growth medium (HuMedia-EG2, Kurabo, Osaka, Japan) supplemented with 2% fetal bovine serum in a 5% CO_2_ incubator (MCO-5AC, Panasonic, Osaka, Japan). When the cultured cells reached 80% confluency in the 96-well plates, incubation continued for an additional 12 h in medium with or without the addition of sample extract. Culture supernatants were collected by centrifugation at 1,000 × *g* for 15 min at room temperature (25 °C), and then reduced with nitrate reductase and the respective enzyme cofactors (iron, molybdenum, and cytochrome) for 30 min at 37 °C. This was followed by 15-min incubation with DAN. Fluorescence intensity was measured at an excitation of 360 nm and emission of 450 nm. The concentration of NO per sample was calculated by transforming raw data using a calibration curve (correlation coefficient R^2^ = 0.9905) prepared with NaNO_3_. The results were expressed as relative values derived from a comparison with the control value of 1.

### Statistical Analysis

Quantitative analysis of the pungent components involved measuring the changes in the contents of pungent components in leaves and fruits over time, measuring antioxidant activity, and NO quantification using a modified Griess method. Each analysis was repeated four times independently. Significant differences among all groups were assessed using Student’s *t*-test and analysis of variance (ANOVA). Data were shown as the mean ± standard deviation (SD). A value of *P* < 0.05 was considered statistically significant.

## Results and Discussion

The capsaicinoid contents of the seven *Satsuma*-Capsicum strains grown in sandy soils were separated into pericarp plus s/p (Figure 1). Calibration curves for capsaicin and dihydrocapsaicin were obtained for each standard. The correlation coefficients were R^2^ > 0.9998 (capsaicin) and R^2^ > 0.9996 (dihydrocapsaicin). Chromatograms showed retention times of 9.1 min for capsaicin and 12.5 min for dihydrocapsaicin. The pungent component contents for the pericarps and s/p per 1 g of each cultivar of Capsicum was quantified as depicted in Figure 2. The s/p from *Satsuma*-Habanero orange, *Satsuma*-Taruna, and *Satsuma*-Tabasco peppers contained higher concentrations of pungent components than the pericarps. Previous studies have reported that pungent components migrate and disperse from the placenta to the pericarp [18–20]. Thus, it is possible that the transition of pungent components in *Satsuma*-Habanero red, *Satsuma*-Indonesian origin, and *Satsuma*-Okinawan chili pepper, and that of Habanero orange in tropical-origin pepper, was faster than that in other cultivars. Furthermore, s/p from Habanero orange and Tabasco pepper *Satsuma*-Capsicum plants contained higher levels of capsaicinoids than those from tropical-origin peppers. On the other hand, their levels in pericarps were similar in both groups. In Habanero orange and Tabasco pepper, pungent components were not detected in any of the leaf or flower samples. Thus, it is likely that Capsicum cultivated under adverse growing conditions contained higher levels of capsaicinoids in the s/p.

**Figure 2.**
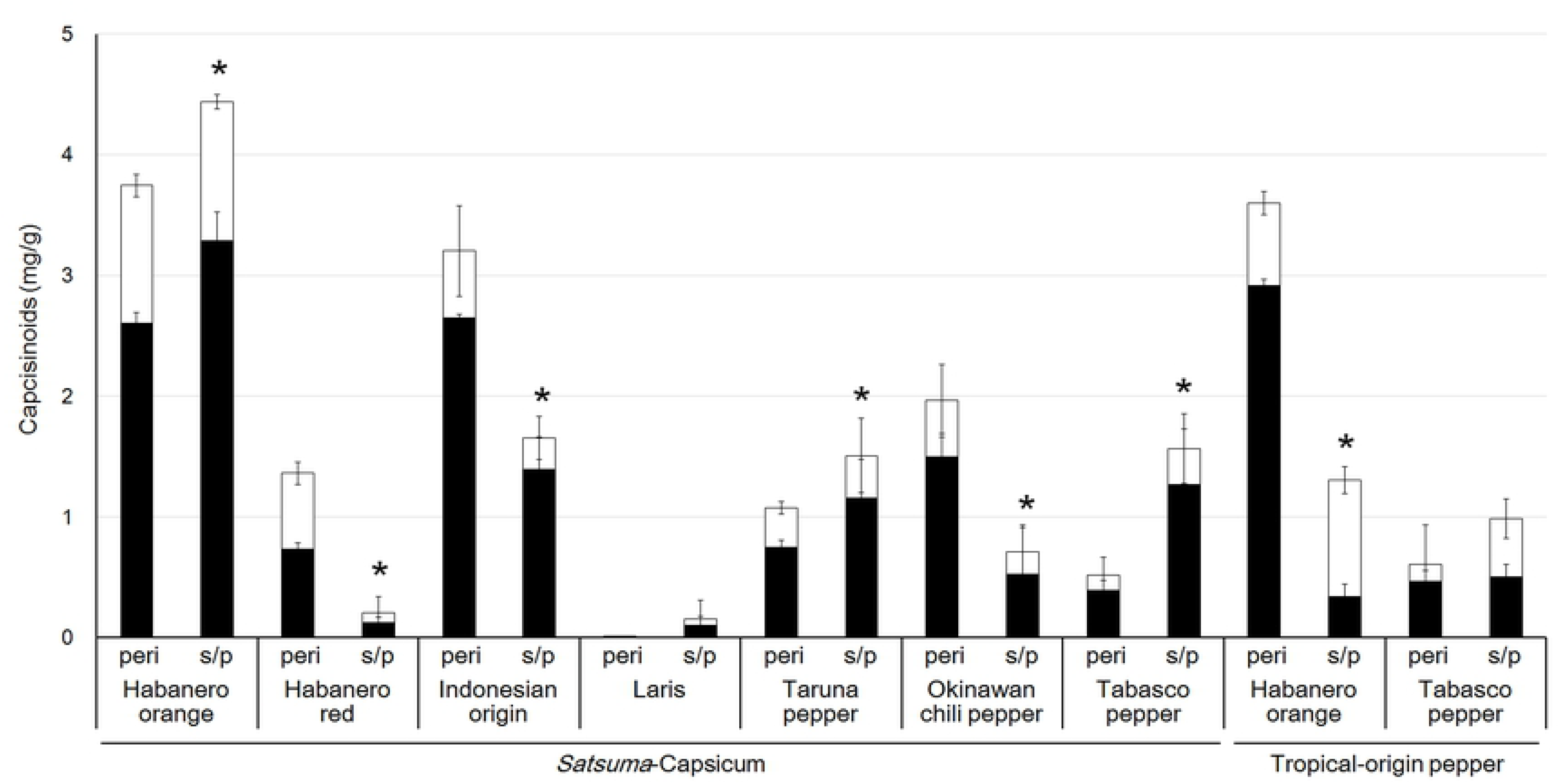
Quantification of capsaicinoids from *Satsuma*-Capsicum and tropical-origin peppers. Black bars represent capsaicin, while white bars represent dihydrocapsaicin. Peri, pericarps; s/p, seeds and placentas. \**P* < 0.05 versus total capsaicinoids in pericarps from the same pepper.

In *Satsuma*-Capsicum, the bottom leaves from Habanero orange (Table 1) and Tabasco pepper (Table 2) were deeper in color and larger than the top leaves. In December, when the temperature rapidly decreased, the leaves shriveled and turned yellow, and the fruit withered. After bearing fruit, the lengths of the leaves exceeded 10 cm. However, pungent components were not detected in any leaf or flower samples. The pungent compounds in the fruit of the Habanero orange reached maximum levels approximately 136 days after planting Habanero orange (approximately 86 days after flowering and fruiting; collection No. 18), and approximately 113 days after planting Tabasco pepper (approximately 63 days after flowering and fruiting; collection No. 21). The peak point varied between cultivars (Figure 3). Even after the leaves yellowed in December, the withered fruit contained abundant levels of pungent components. We did not identify any correlations between the pungent component contents in the fruit and the color of the leaves and fruit. In order to collect the fruits at an optimal time, it was necessary to consider the time between planting, flowering, and fruiting, rather than simply observing visible changes. Furthermore, *Satsuma*-Capsicum was slower to wither than tropical-origin peppers and could be harvested for a longer time period in our study.

**Table 1.**
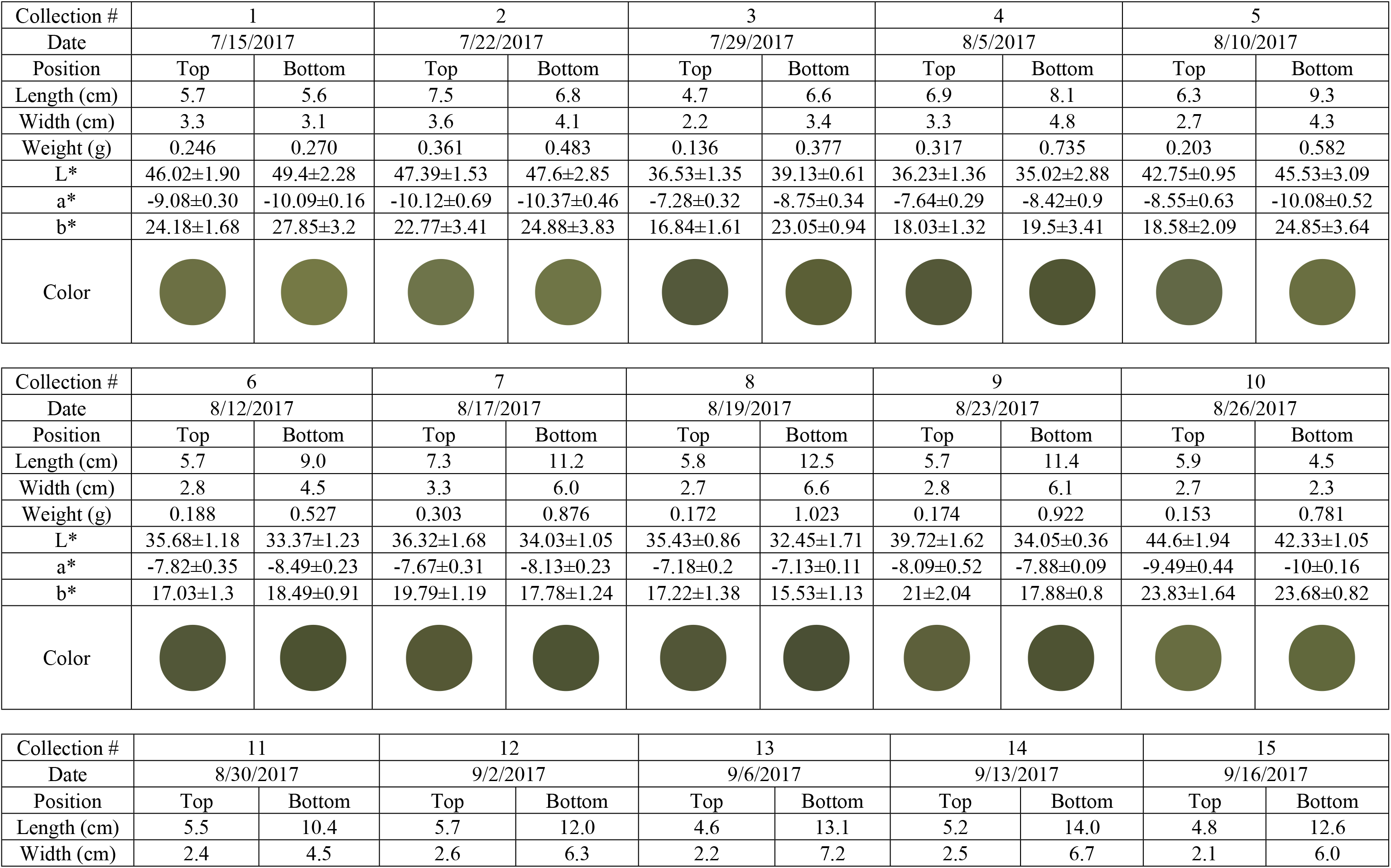
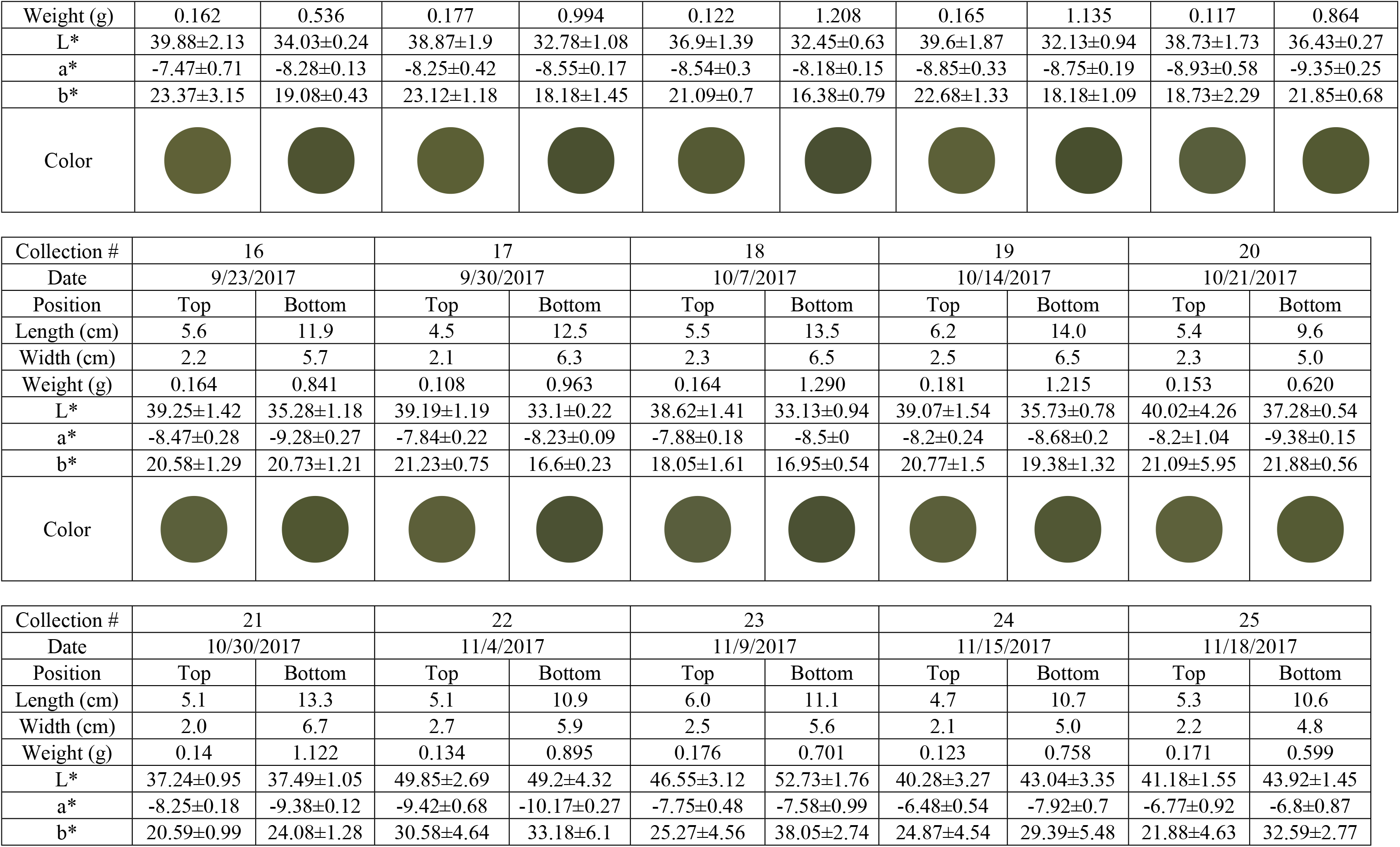
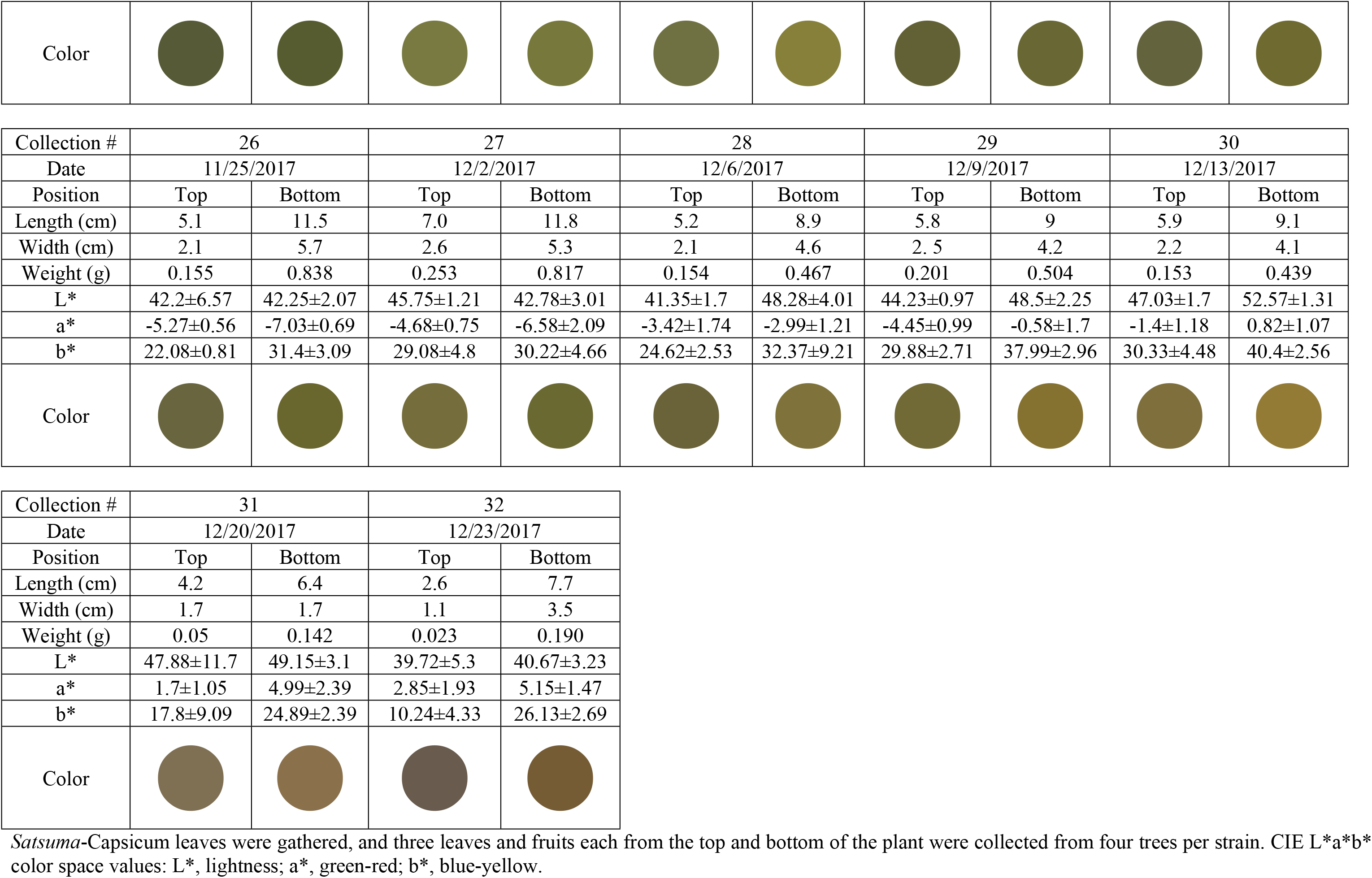
Leaf characteristics during the growth period of the *Satsuma*-Habanero orange pepper.

**Table 2.**
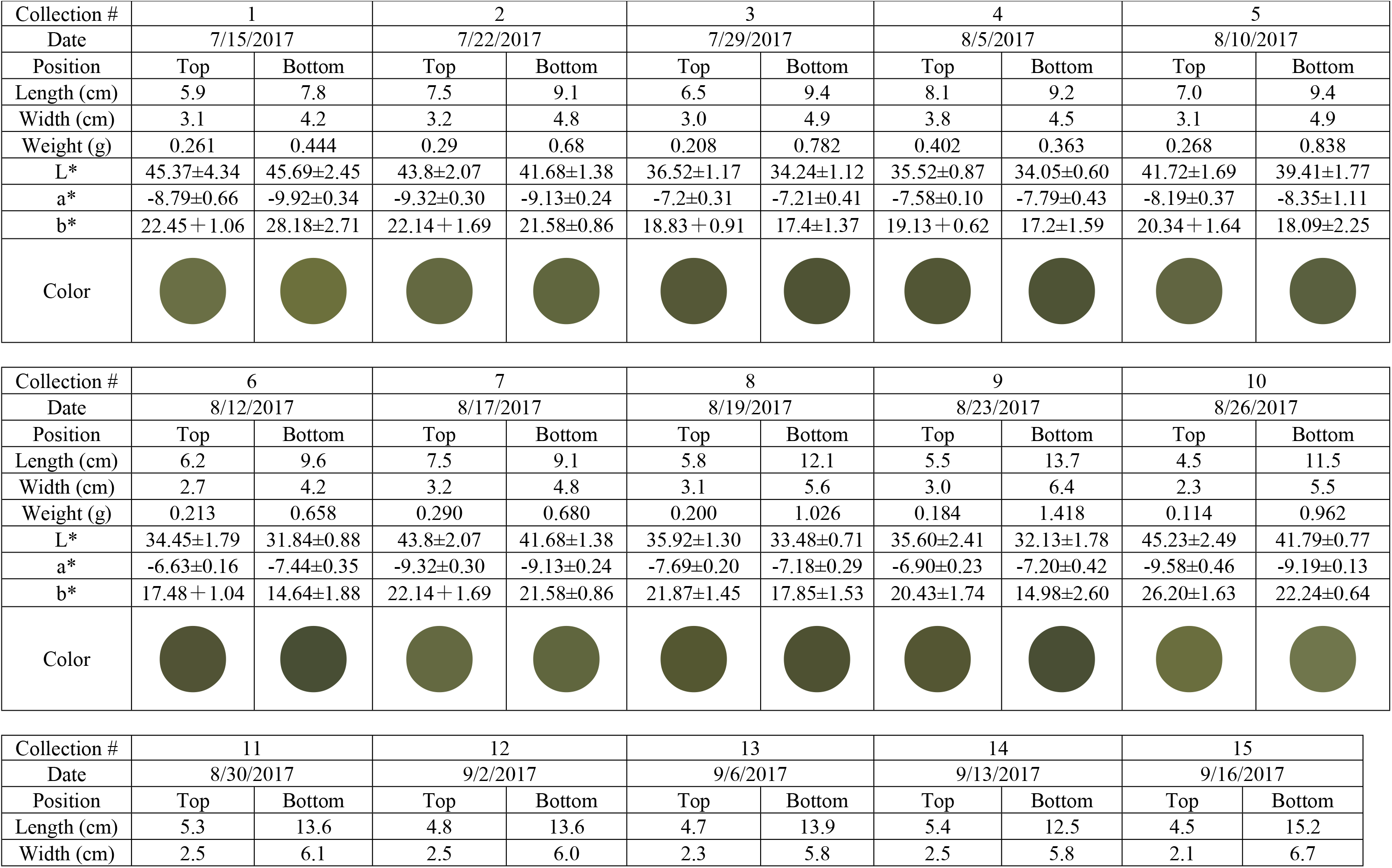
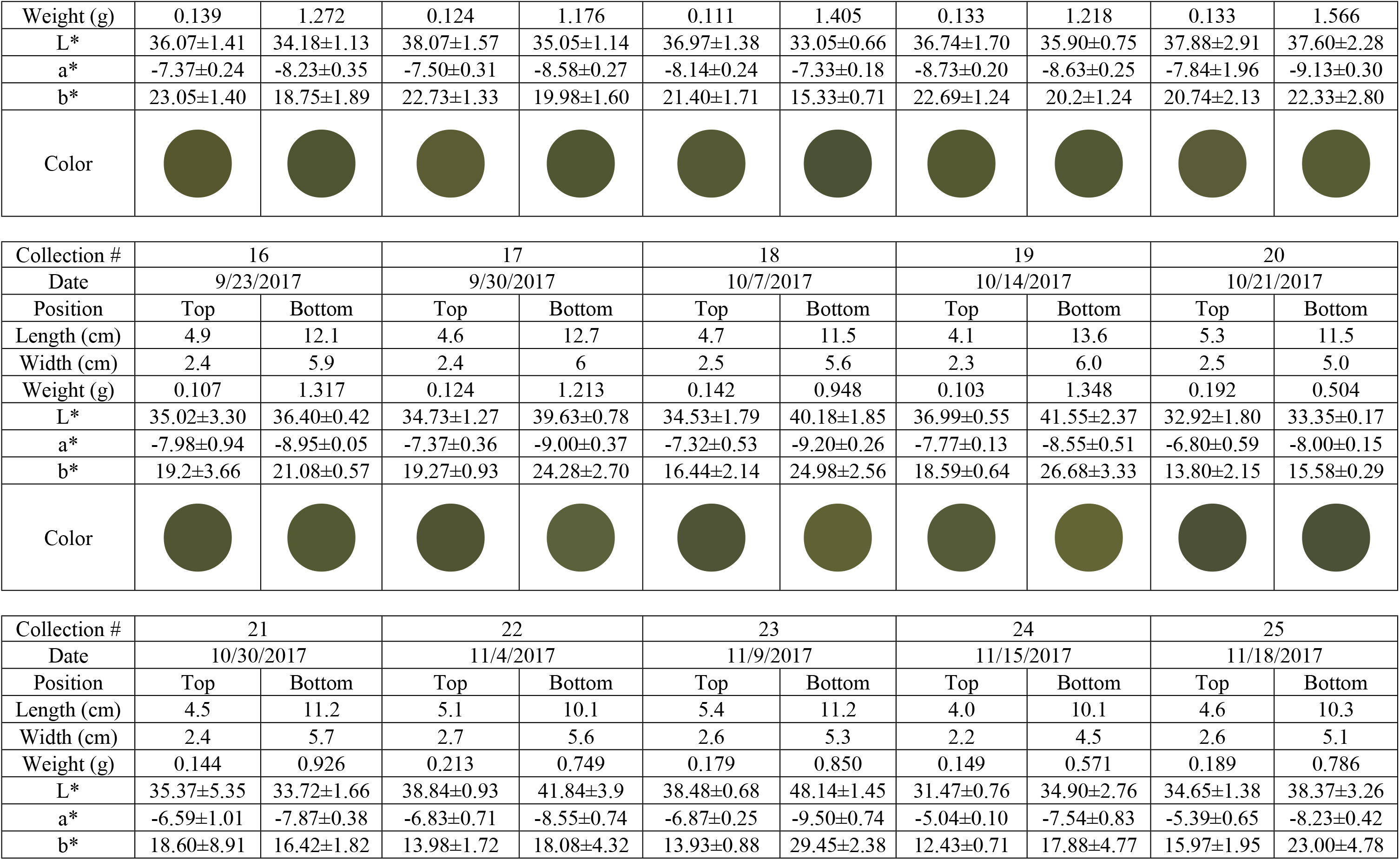
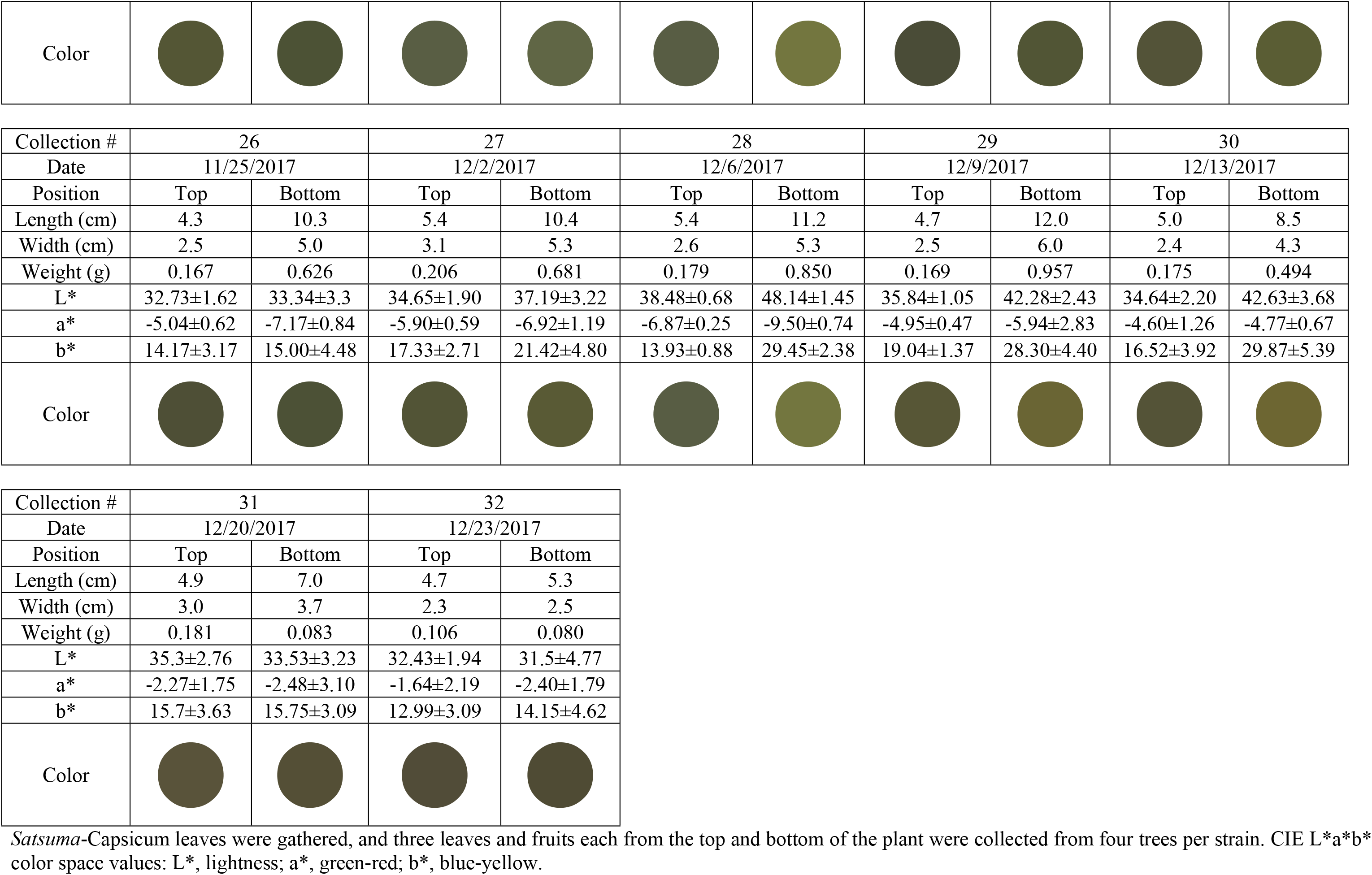
Leaf characteristics during the growth period of the *Satsuma*-Tabasco pepper.

**Figure 3.**
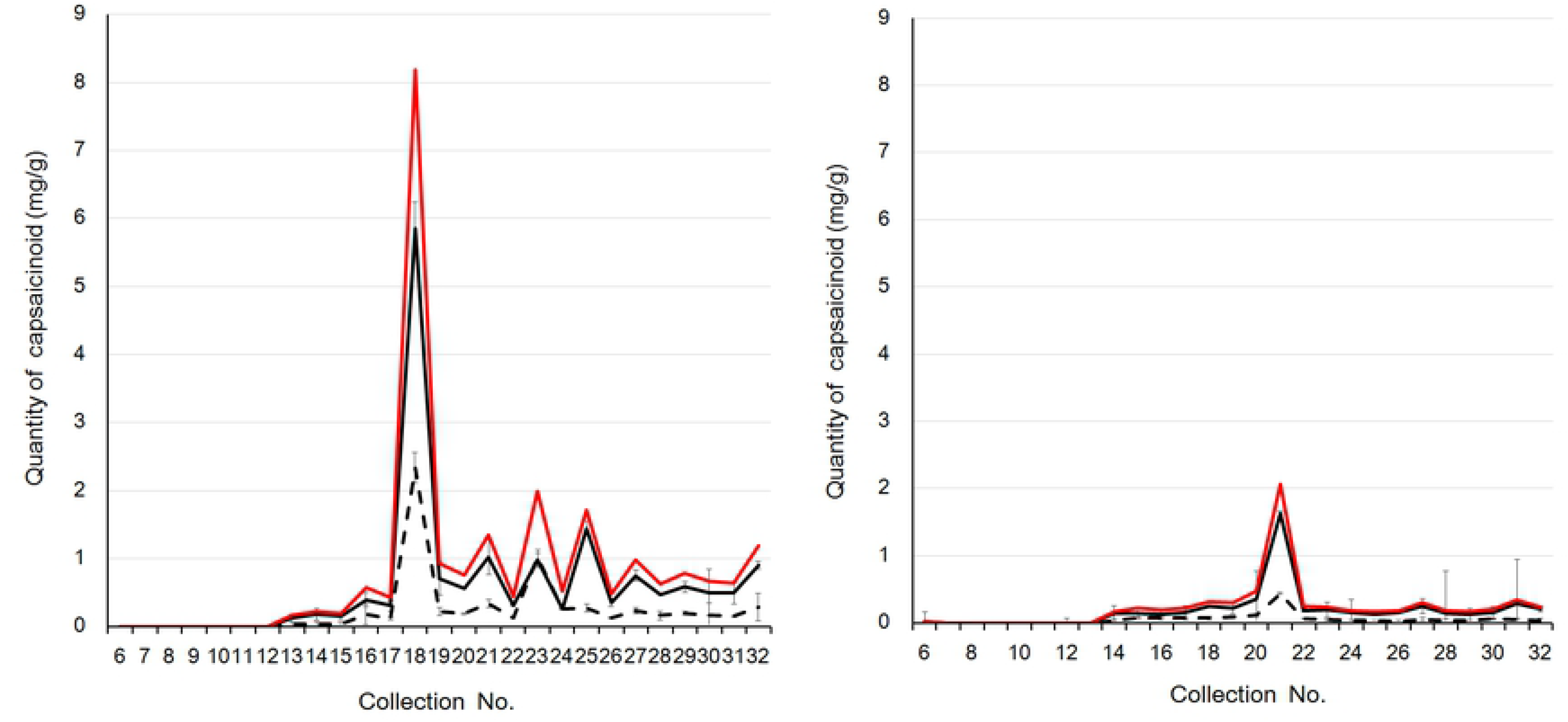
Changes over time in the concentration of capsaicin and dihydrocapsaicin in (A) *Satsuma*-Habanero orange and (B) *Satsuma*-Tabasco pepper fruits. Solid black line, capsaicin; black dotted-line, dihydrocapsaicin; solid red line, total capsaicin and dihydrocapsaicin contents.

All *Satsuma*-Capsicum plants induced antioxidative activity *in vitro* (Figure 4A). The antioxidative capacity of *Satsuma*-Habanero orange was significantly higher than that of tropical-origin Habanero orange in the s/p, but not in the pericarps. In contrast, antioxidative activity was significantly higher in the pericarp of Habanero red and Indonesian-origin peppers than that in the s/p. There was no correlation between antioxidant activity and pungent compound content s (R^2^ = 0.4579). Capsicum consists of many functional components such as carotenoid pigments, capsanthin, and α-tocopherol, in addition to the pungent ingredients. Thus, we concluded that the antioxidant properties might be due to interactions between the liposoluble carotenoid pigments capsanthin and α-tocopherol, rather than due to the pungent components [21,22].

**Figure 4.**
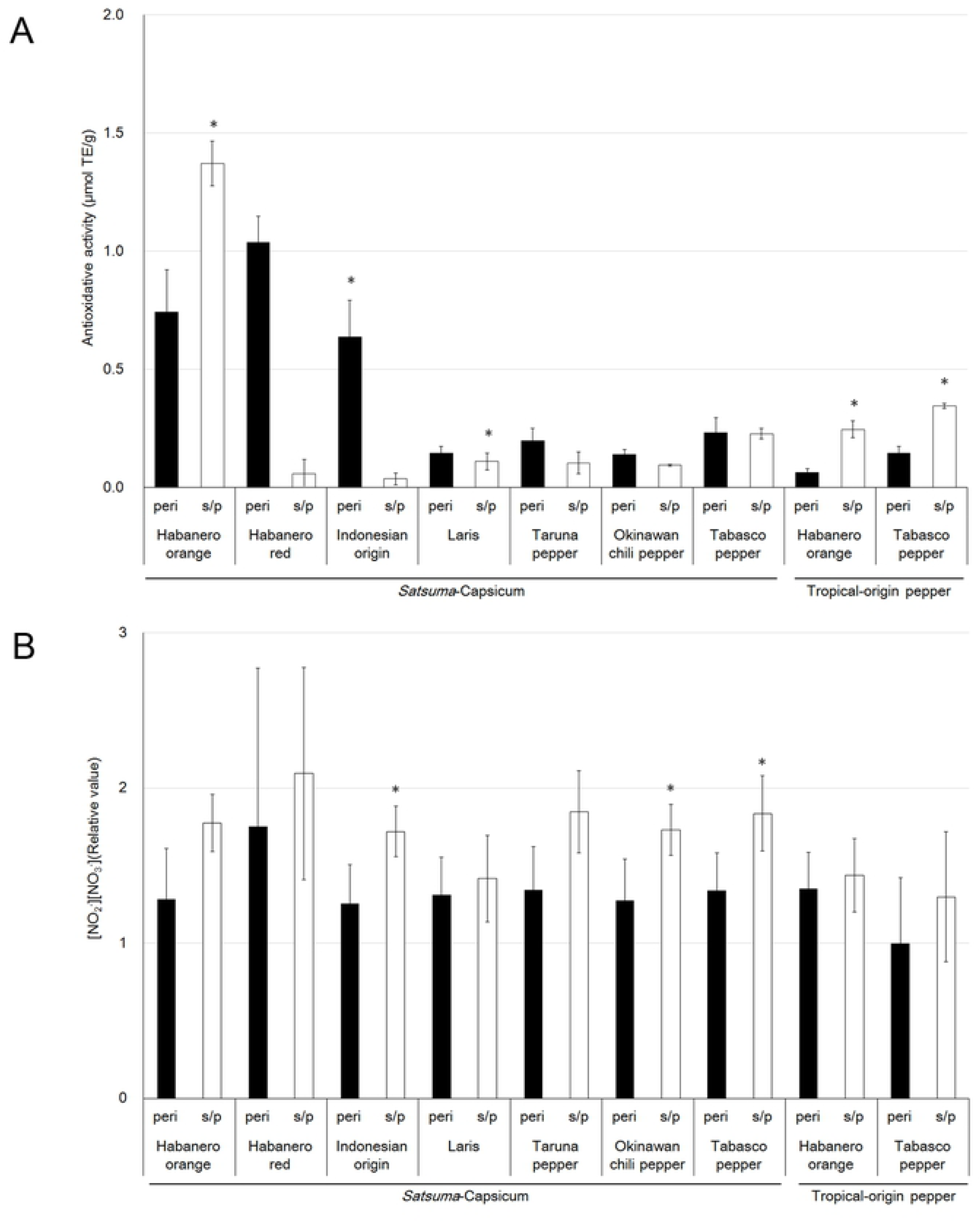
(A) Antioxidative activity and (B) the level of nitric oxide (NO) production in vascular endothelial cells for *Satsuma*-Capsicum and tropical-origin peppers. Peri, pericarps; s/p, seeds and placentas. \**P* < 0.05 versus pericarps

We further investigated whether *Satsuma*-Capsicum would promote NO production in VECs at levels similar to those of tropical-origin peppers to confirm their bioregulatory effect on vascular function. Our results indicated that all peppers could potentially increase NO production, but there was no significant difference between pericarps and s/p. Additionally, these data also showed that the effects of *Satsuma*-Capsicum extracts on NO production were similar to the effects of the tropical-origin pepper extracts (Figure 4B).

## Conclusions

All *Satsuma*-Capsicum plants cultivated under adverse conditions in Southern Japan showed higher antioxidative activity than the traditionally grown tropical peppers. Therefore, *Satsuma*-Capsicum extracts from peppers cultivated in harsh environments within Southern Japan showed similar effects on NO production to those of tropical-origin pepper extracts; thus, *Satsuma*-Capsicum can potentially improve vascular endothelial function. We also concluded that Capsicum cultivated under these adverse conditions contained higher levels of capsaicinoids than those cultivated in tropical regions. Moreover, *Satsuma*-Capsicum plants cultivated in nutrient-poor sandy soil contained higher concentrations of pungent components than traditional tropical-origin peppers. Finally, the *Satsuma*-Capsicum plants cultivated in this study were slower to spoil than traditional tropical-origin peppers and could be harvested over a longer period of time.

Capsaicin and dihydrocapsaicin are pungent components within peppers. The continuous consumption of capsaicin has been found to promote fat reduction in humans [23,24], the induction of skeletal muscle hypertrophy [25], and has shown potential for reducing obesity [26,27]. These effects could be the result of activity by capsaicin, a vanilloid belonging to the vanillyl group. Capsaicin potentially stimulates the transient receptor potential cation channel subfamily V member 1 (TRPV1), a receptor activation channel, by binding to vanilloid receptors; this could promote lipolysis and generate heat [28–31].

Other varieties of pepper and paprika are also widely considered to promote health, as they exhibit high antioxidant activity and also possess properties shown to limit the proliferation of cancer cells [32–34]. Because paprika contains only trace amounts of capsaicin and dihydrocapsaicin, these properties were thought to result from the activity of capsanthin or carotenoid pigments. Moreover, peppers have been reported to contain polyphenols with various biomodulation functions [35–37]. The *Satsuma*-Capsicum plants investigated in this study showed significant antioxidative activity, which may have resulted from polyphenols, as there was no correlation between antioxidant activity and the concentration of pigment (R^2^ = 0.4579).

Our results indicated that *Satsuma*-Capsicum plants are superior to traditional tropical-origin peppers, and thus show greater value for future product development.

## Acknowledgements

The authors would like to thank Dr. Fumio Yagi for advice with the experiments and thank Dr. Jun-Ichi Sakagami and Mr. Kenta Komori for providing Capsicum samples.

